# TMEM16F phospholipid scramblase mediates trophoblast fusion and placental development

**DOI:** 10.1101/711473

**Authors:** Yang Zhang, Trieu Le, Ryan Grabau, Zahra Mohseni, Hoejeong Kim, David R. Natale, Liping Feng, Hua Pan, Huanghe Yang

## Abstract

Cell-cell fusion or syncytialization is fundamental to the reproduction, development and homeostasis of multicellular organisms. In addition to various cell-type specific fusogenic proteins, cell surface externalization of phosphatidylserine (PS), a universal eat-me signal in apoptotic cells, has been observed in different cell-fusion events. Nevertheless, molecular underpinnings of PS externalization and cellular mechanisms of PS-facilitated cell-cell fusion are unclear. Here we report that TMEM16F, a Ca^2+^-activated phospholipid scramblase (CaPLSase), plays an indispensable role in placental trophoblast fusion by translocating PS to the cell surface independent of apoptosis. Consistent with its essential role in trophoblast fusion, the placentas from TMEM16F-deficient mice exhibit deficiency in syncytialization, placental developmental defects and perinatal lethality. Our findings thus identify a cell-cell fusion mechanism by which TMEM16F CaPLSase-dependent externalization of PS serves as a critical cell fusion signal to facilitate trophoblast syncytialization and placental development.

---

Cell-cell fusion is a fundamental cellular process essential for sexual reproduction, development and homeostasis in organisms ranging from fungi to human^1-3^. Sperm-egg fusion determines the successfulness of fertilization^4^. Myoblast fusion into multinucleated myofibers is required for the growth, maintenance and regeneration of skeletal muscles^5^. Fusion of macrophages into bone-absorbing osteoclasts and giant cells is critical for bone homeostasis and immune response, respectively^6^. In the human and mouse placentas, mononucleated cytotrophoblasts continuously fuse with overlying syncytiotrophoblasts to sustain barrier function and transport activities^7^. Consistent with its critical physiological roles, malfunction of cell-cell fusion has been implicated in various of diseases ranging from infertility, myopathies, osteoporosis, cancer and preeclampsia^1, 3, 8, 9^.

Despite its importance in biology and pathology, cell-cell fusion remains poorly understood compared to other membrane fusion processes^3^. The recent progresses in cell-cell fusion mechanisms have successfully identified a number of fusogenic proteins or fusogens in mammals, such as myomaker (TMEM8C) and myomixer (Gm7325/Minion) that are specifically expressed in vertebrate myoblasts^10, 11^, as well as syncytin proteins of retroviral origin that are solely expressed in placental trophoblasts^12,13^. The identifications of the fusogens greatly advanced our understanding the mechanisms of cell-cell fusion. However, cell-type specific expression of these fusogens implies that cell-cell fusion is a highly diverse and specialized cellular process^1-3^.

As an anionic phospholipid that predominantly resides in the inner leaflet of the plasma membrane of all eukaryotic cells, surface exposure of PS is known to play multifaceted roles including attracting blood clotting factors to enable blood coagulation and serving as an eat-me signal to attract phagocytes^14-16^. Interestingly, PS surface exposure has also been observed in various cell-cell fusion events such as myoblast fusion, sperm-egg fusion and trophoblast fusion^3, 14^. Compared to cell-type specific fusogens, PS exposure seems to be a more generalized feature of cell-cell fusion of different cell types. The notion of PS exposure as an early hallmark of apoptosis led to the prevailing view of PS exposure during cell-cell fusion: apoptosis is responsible for PS exposure-mediated cell-cell fusion^17-19^. Nevertheless, syncytialization determines the reproduction and survival of a species. Fusing apoptotic cells to the physiologically important multi-nucleated cells seems rather harmful than beneficial to the fused cells. It is thus critical to understand the molecular underpinning of PS exposure during cell fusion and the mechanism on how PS exposure mediates cell-cell fusion.

Here we report that TMEM16F, a Ca^2+^-activated phospholipid scramblase (CaPLSase) that translocates PS to platelet surface and facilitates blood coagulation^20, 21^, mediates PS exposure during trophoblast fusion independent of apoptosis. Genetic ablation of TMEM16F abolishes fusion among BeWo cells, a human choriocarcinoma trophoblast cell line, while reintroducing TMEM16F back to the knockout cells rescues BeWo cell fusion. Consistent with the *in vitro* findings, the placentas of TMEM16F-deficient mice exhibit trophoblast fusion deficiency, developmental defects in the labyrinth layer and fetal blood vessels, resulting in perinatal lethality. Our findings thus uncover a cell-cell fusion mechanism that requires CaPLSase-mediated PS exposure as an important cell fusion signal. The CaPLSase-mediated cell-cell fusion mechanism can shine light on understanding other cell fusion events as well as cell fusion in health and disease.

## Results

### PS surface externalization is required for trophoblast fusion

Forskolin-induced BeWo cell fusion is widely used to study the cellular mechanism of villous trophoblast fusion *in vitro*^22, 23^. Using Di-8-ANEPPS dye to mark the plasma membrane (Figure 1A), the fusion index of live BeWo cells, which is defined as the percentage of nuclei from cells with ≥ 3 nuclei (to exclude potential dividing cells, see Methods for details), can be reliably quantified^24^. 30 µM forskolin stimulation significantly boosts the fusion index from 0.03±0.01 to 0.38±0.02 (Figure 1B and 1E); whereas application of Annexin V (AnV), a specific PS-binding protein, almost completely suppresses forskolin-induced BeWo cell fusion (Figure 1C and 1E). Our results support that PS surface exposure plays a critical role in trophoblast fusion which is consistent with the previous reports showing that masking surface-exposed PS with PS-specific monoclonal antibodies inhibits BeWo cell fusion^25,26^.

**Figure 1.**
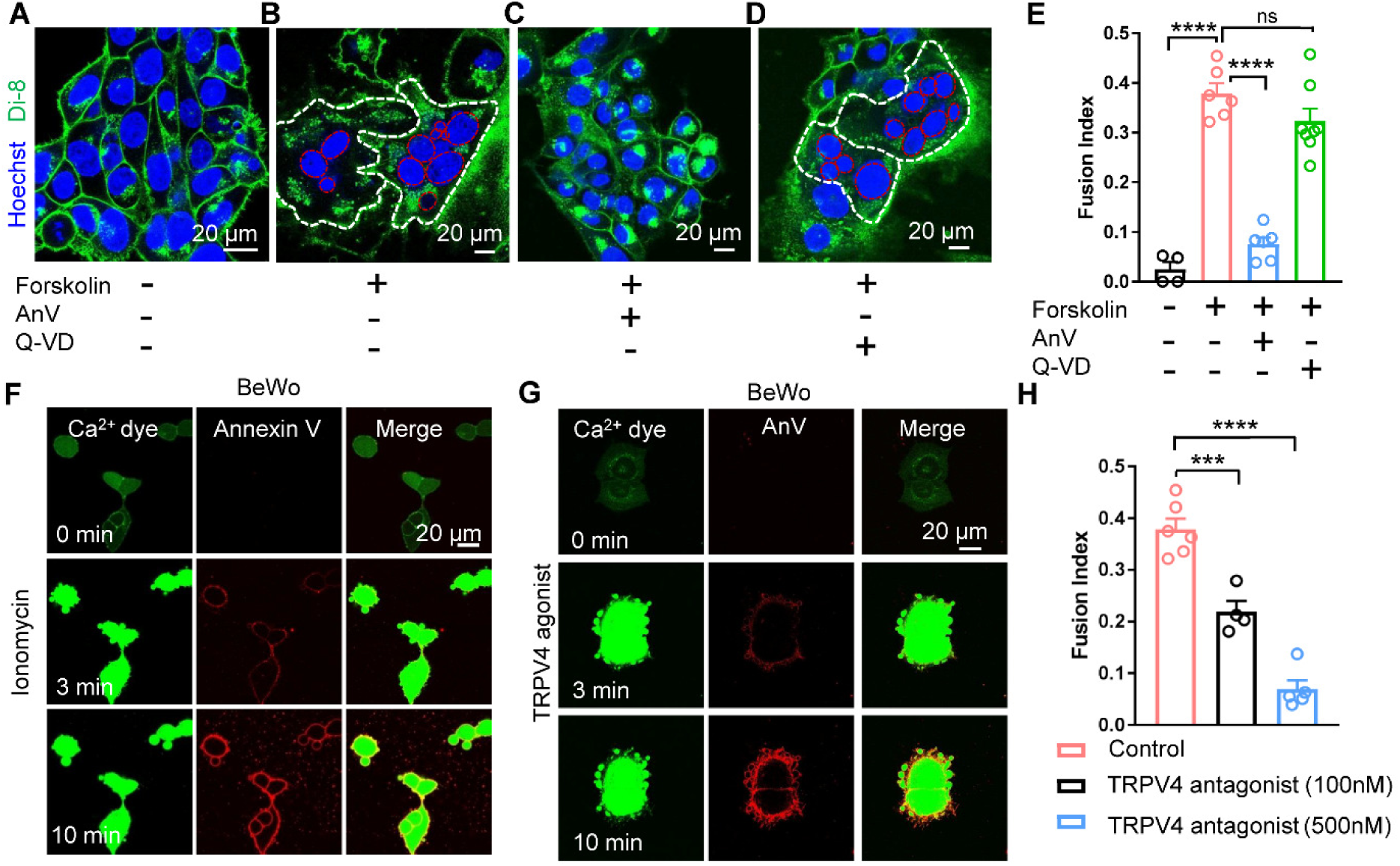
Ca^2+^-activated lipid scrambling is required for trophoblast fusion. (A) Di-8-ANEPPS (Di-8), a membrane-bound voltage sensitive dye, delineates BeWo trophoblast boundaries (green). Hoechst label nuclei (blue). (B) 30 µM forskolin induces BeWo trophoblast fusion in 48 h. White and red dotted lines delineate the plasma membrane and the nuclei of the fused cells, respectively. (C) 0.5 µg/ml Annexin V (AnV), a PS binding protein, inhibits forskolin-induced BeWo cell fusion. (D) 10 µM pan-caspase inhibitor Q-VD-OPH (Q-VD) does not affect forskolin-induced BeWo cell fusion. (E) Summary of forskolin, AnV and Q-VD effects on BeWo cell fusion. (F, G) BeWo cells exhibit robust CaPLSase activities when triggered with 1 µM ionomycin (F) or 20 nM GSK1016790A (G), a potent TRPV4 channel agonist (see also Movies S1 and S2). Ca^2+^ dye (Calbryte 520) and fluorescently tagged AnV proteins (AnV-CF594) are used to measure the dynamics of intracellular Ca^2+^ and PS externalization, respectively. (H) Forskolin-induced BeWo cell fusion is inhibited by GSK2193874, a selective TRPV4 Ca^2+^-permeable channel antagonist. Unpaired two-sided Student’s *t*-test for panels E and H. ***p < 0.001, ****p < 0.0001. ns: not significant. Error bars indicate ± SEM. Each dot represents the average of fusion indexes of six random fields from one coverslip (see Methods for detailed quantification). All fluorescence images are the representatives of at least three biological replicates.

### Ca^2+^-activated but not caspase-activated phospholipid scramblases are critical for trophoblast fusion

Phospholipid scramblases are passive phospholipid transporters on cell membranes that catalyze PS surface exposure^14^. There are two major types of phospholipid scramblases. Caspase-dependent lipid scramblases such as Xkr8 induce PS externalization^27^, which serves as an eat-me signal in apoptotic cells to attract phagocytes and initiate phagocytosis^16^. CaPLSases, on the other hand, mediate rapid PS surface exposure in viable cells in response to intracellular Ca^2+^ elevation^14, 16, 20^. To pinpoint the specific type of phospholipid scramblases that are responsible for PS externalization in trophoblast-derived BeWo cells, we first treated the cells with Q-VD-OPh, a pan-caspase inhibitor. The lack of inhibition on the fusion index by Q-VD-OPh (Figure 1D and 1E) indicates that that caspase-dependent lipid scrambling is not required for BeWo cell fusion. Our result is consistent with previous findings, which showed that apoptosis does not make major contributions to trophoblast fusion^28, 29^.

Instead, the human trophoblasts exhibited robust CaPLSase activity measured by an optimized phospholipid scrambling assay (Figure S1A)^30^. We stimulated both BeWo and primary human term placental trophoblasts with ionomycin, a Ca^2+^ ionophore (Figures 1F, S1B and Movies S1), or GSK-1016790A^31^, a potent and specific agonist of TRPV4 Ca^2+^-permeable channel (Figures 1G, S1D and Movie S2). Upon stimulation, fluorescently-tagged AnV rapidly accumulates on trophoblast surface following intracellular Ca^2+^ elevation (Figure S1A), indicating that functional CaPLSases are abundantly expressed in human trophoblasts. Our results also suggest that TRPV4, a previously unreported Ca^2+^-permeable channel in trophoblasts (Figure S1C), can serve as an upstream Ca^2+^-importer for CaPLSase activation.

Consistent with the observed dominant CaPLSase activity in trophoblasts (Figure 1F and 1G) and the importance of PS externalization in trophoblast fusion (Figure 1E), disruption of Ca^2+^ homeostasis by reducing extracellular Ca^2+^ (Figure S1E) or pharmacologically inhibiting TRPV4 channels with GSK-2193874^31^ hinders BeWo cell fusion (Figure 1H). Therefore, our results demonstrate that CaPLSase-mediated PS exposure plays an essential role in trophoblast fusion.

### TMEM16F Ca^2+^-activated lipid scramblase is highly expressed in human trophoblasts

All currently known CaPLSases are TMEM16 family members^20, 32-34^. To pinpoint the molecular identity of CaPLSase in trophoblast, we measured the relative mRNA levels of the TMEM16 family in both BeWo (Figure 2A) and primary human term trophoblast cells (Figure S2A) by quantitative RT-PCR. Of these, TMEM16F, a CaPLSase critical for platelet PS externalization during blood coagulation^20, 21^, shows the highest expression level in BeWo and human primary trophoblasts, whereas the expression levels of other TMEM16F members are negligible. Immunofluorescence staining confirms that TMEM16F protein is highly expressed in both BeWo and human primary trophoblasts (Figure 2B). In human first trimester and term placentas, TMEM16F is highly expressed in both syncytiotrophoblasts and cytotrophoblasts in the chorionic villi (Figures 2C and S2B). Our results thus demonstrate that TMEM16F is highly expressed in human trophoblasts and is the major CaPLSase responsible for PS surface exposure.

**Figure 2.**
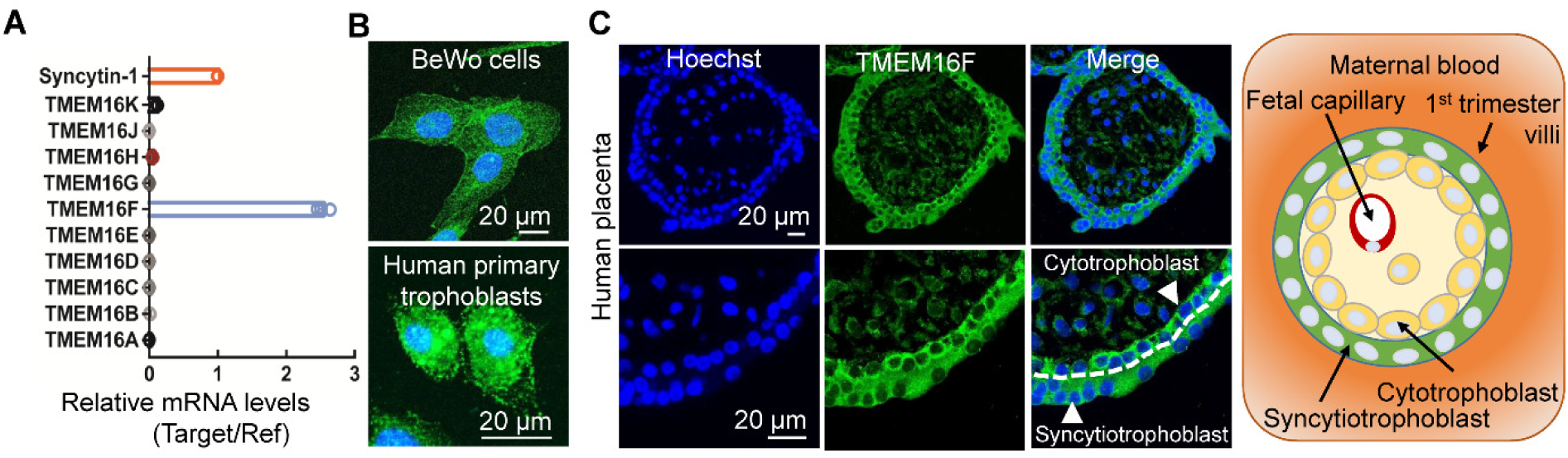
TMEM16F CaPLSase is highly expressed in human placental trophoblasts. (**A**) qRT-PCR of TMEM16 family members in BeWo cells. All genes are normalized to GAPDH and then normalized to Syncytin-1. (**B**) Immunofluorescence of TMEM16F (green) in BeWo cells (upper) and the primary human placental trophoblasts from a term placenta (lower), The nuclei are stained with Hoechst (blue). (**C**) Immunofluorescence of TMEM16F (anti-TMEM16F, green) and nuclei (Hoechst, blue) in a human first trimester placenta villi (upper). Higher magnifications are shown in the lower panels. TMEM16F is highly expressed in both cytotrophoblasts and syncytiotrophoblasts (arrowhead). The white dotted line demarcates the basal membrane of the syncytiotrophoblast. Schematic of the maternal-fetal interface in the first trimester placental villi (right). Cytotrophoblasts tightly pack against the basal membrane of syncytiotrophoblasts, which form the barrier between maternal blood (orange) and fetal capillaries located in the villous stroma. Images and diagram are shown in cross-sections of the villi.

### TMEM16F-CaPLSase is indispensable for trophoblast fusion

To further examine TMEM16F function in trophoblast fusion, we generated a TMEM16F knockout (KO) BeWo cell line using CRISPR/Cas9 methods. Genetic ablation of TMEM16F completely eliminates the protein expression in BeWo cells as examined by western blotting and immunostaining (Figure S3A and S3B). Distinct from the Cas9 control cells that show strong CaPLSase activity (Figure S3C and S3D), the TMEM16F KO BeWo cells lack Ca^2+^-induced PS exposure when stimulated with ionomycin (Figure 3A and Movie S3) and TRPV4 agonist GSK-1016790A (Figure S3E). The intracellular Ca^2+^ elevation in response to ionomycin and GSK-1016790A remain intact (Figures 3A and S3E, Movies S3 and S4), indicating that the deficiency in phospholipid scrambling in the TMEM16F KO BeWo cells is due to the lack of functional CaPLSases but not impaired Ca^2+^ response.

**Figure 3.**
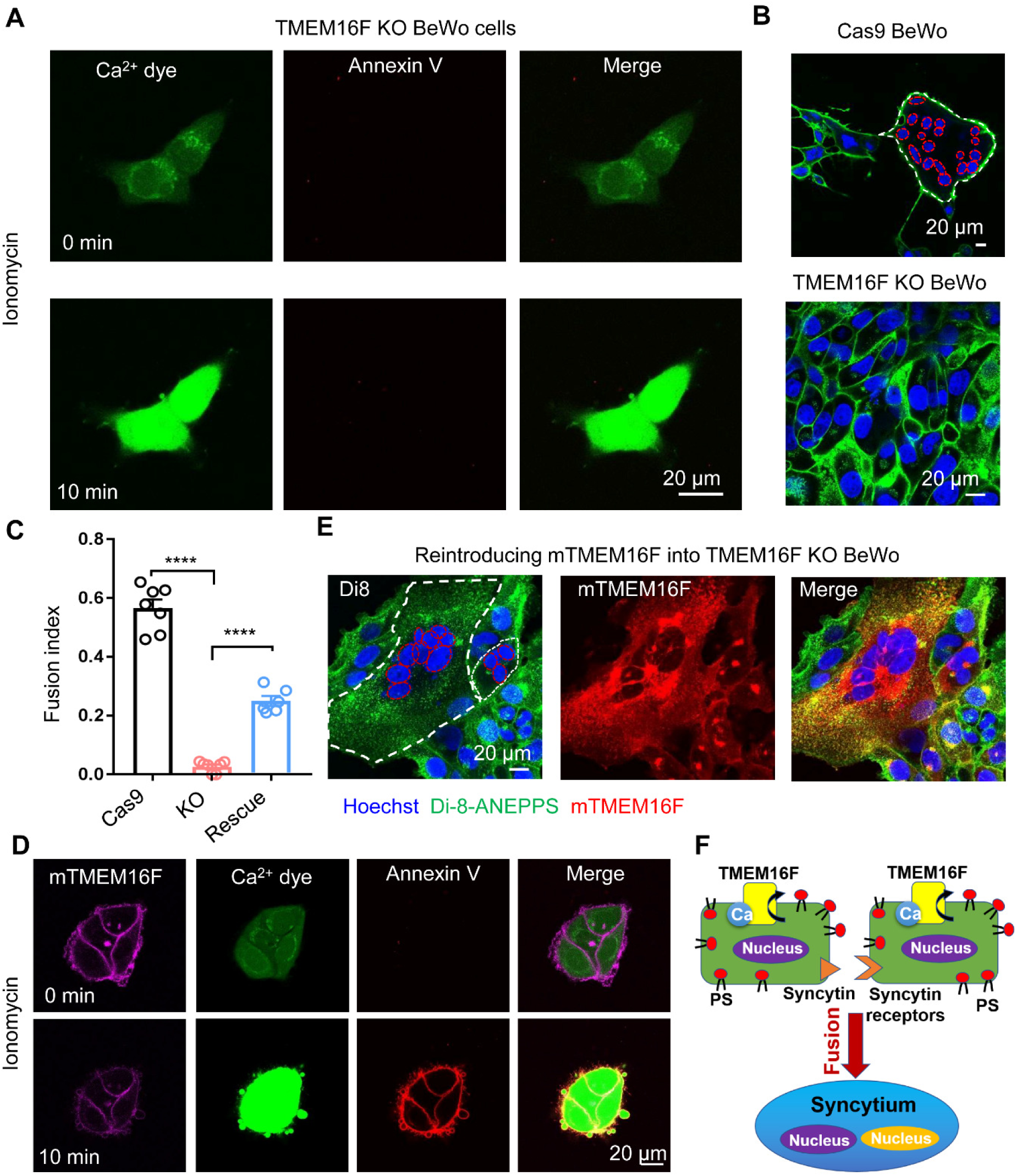
TMEM16F CaPLSase is indispensable for trophoblast fusion. (A) CaPLSase activity is eliminated in TMEM16F KO BeWo cells when triggered with 1 µM ionomycin (see also Movies S3). (B) Representative images of Cas9 control cells (upper) and TMEM16F KO BeWo cells (lower) after 48 h forskolin treatment. (C) Genetic ablation of TMEM16F CaPLSase inhibits forskolin-induced BeWo cell fusion and reintroducing mTMEM16F partially rescues the fusion deficiency. Each dot represents the average of fusion indexes of six random fields from one coverslip. Unpaired two-sided Student’s *t*-test. ****p < 0.0001. Error bars indicate ± SEM. (D) Heterologous expression of murine TMEM16F (mTMEM16F) in the TMEM16F KO BeWo cells re-introduces CaPLSase activity (see also Movies S4). 1 µM ionomycin was used to stimulate mTMEM16F. (E) Representative images of the TMEM16F KO BeWo cells overexpressing mTMEM16F after 48 h forskolin treatment. (F) A cell-cell fusion mechanism requires CaPLSase-induced PS externalization on cell surface. All fluorescence images are the representatives of at least three biological replicates. Nuclei and membranes are labelled with Hoechst (blue) and Di-8-ANEPPS (green), respectively in panel B and E. White and red dotted lines delineate the plasma membrane and the nuclei of the fused cells, respectively.

Consistent with the critical role of PS externalization in BeWo cell fusion (Figure 1C), the TMEM16F deficient BeWo cells lacking CaPLSase activity (Figure 3A) fail to undergo fusion after forskolin stimulation (Figure 3B and 3C). Another independent TMEM16F KO BeWo cell line which was generated using a different sgRNA also shows the same deficiency in CaPLSase activity and cell fusion was observed (Data not shown), ruling out potential off-target effects of CRISPR/Cas9 genome engineering. In addition, we heterologously expressed murine TMEM16F (mTMEM16F) to the TMEM16F deficient BeWo cells. Re-introducing mTMEM16F not only restores their CaPLSase activity (Figure 3D and Movie S4) but also rescues cell-cell fusion (Figure 3C and 3E). Taken all together, our TMEM16F ablation and rescue experiments *in vitro* explicitly demonstrate that TMEM16F CaPLSase plays an indispensable role in BeWo trophoblast fusion. It is likely TMEM16F CaPLSase-mediated PS exposure works in concert with trophoblast-specific fusogenic proteins such as syncytins and their receptors to enable trophoblast fusion (Figure 3F).

### TMEM16F knockout mice exhibit deficiency on trophoblast fusion, placental development and perinatal viability

To understand the function of TMEM16F CaPLSase in trophoblast physiology and placental development *in vivo*, we examined the pregnant mice from a TMEM16F deficient (KO) line^21^. In the C57BL/6 genetic background, heterozygous x heterozygous breeding scheme shows a significant reduction of postnatal homozygous TMEM16F^-/-^mice (14.7%) from the expected Mendelian inheritance (25%) (Figure 4A and Table S1). Furthermore, the weights of TMEM16F^-/-^placentas and embryos are significantly reduced compared to wildtype (WT, TMEM16F^+/+^) placentas (Figure 4B and 4C). Detailed histological analysis confirms the placental defects in the TMEM16F^-/-^mice. The KO placentas look apparently pale from the fetal surface and exhibit markedly decreased and unevenly distributed vascularization in both placental and extra-embryonic membranes (Figure 4D and 4E). Consistent with these gross morphological changes, hematoxylin and eosin (H&E) staining of KO placentas reveal markedly enlarged cavities in the labyrinth layer, where syncytiotrophoblasts reside (Figure 4F (i and ii) and 4G (i and ii)). The enlarged cavities are concentrated proximal to the fetal interface. Using alkaline phosphatase (AP) staining^35^ to assess for the sinusoidal trophoblast giant cells (STGCs), we confirmed that the enlarged cavity in the KO placentas are maternal blood sinuses, which is enclosed by STGCs (Figure 4F(iv) and 4G(iv)).

**Figure 4.**
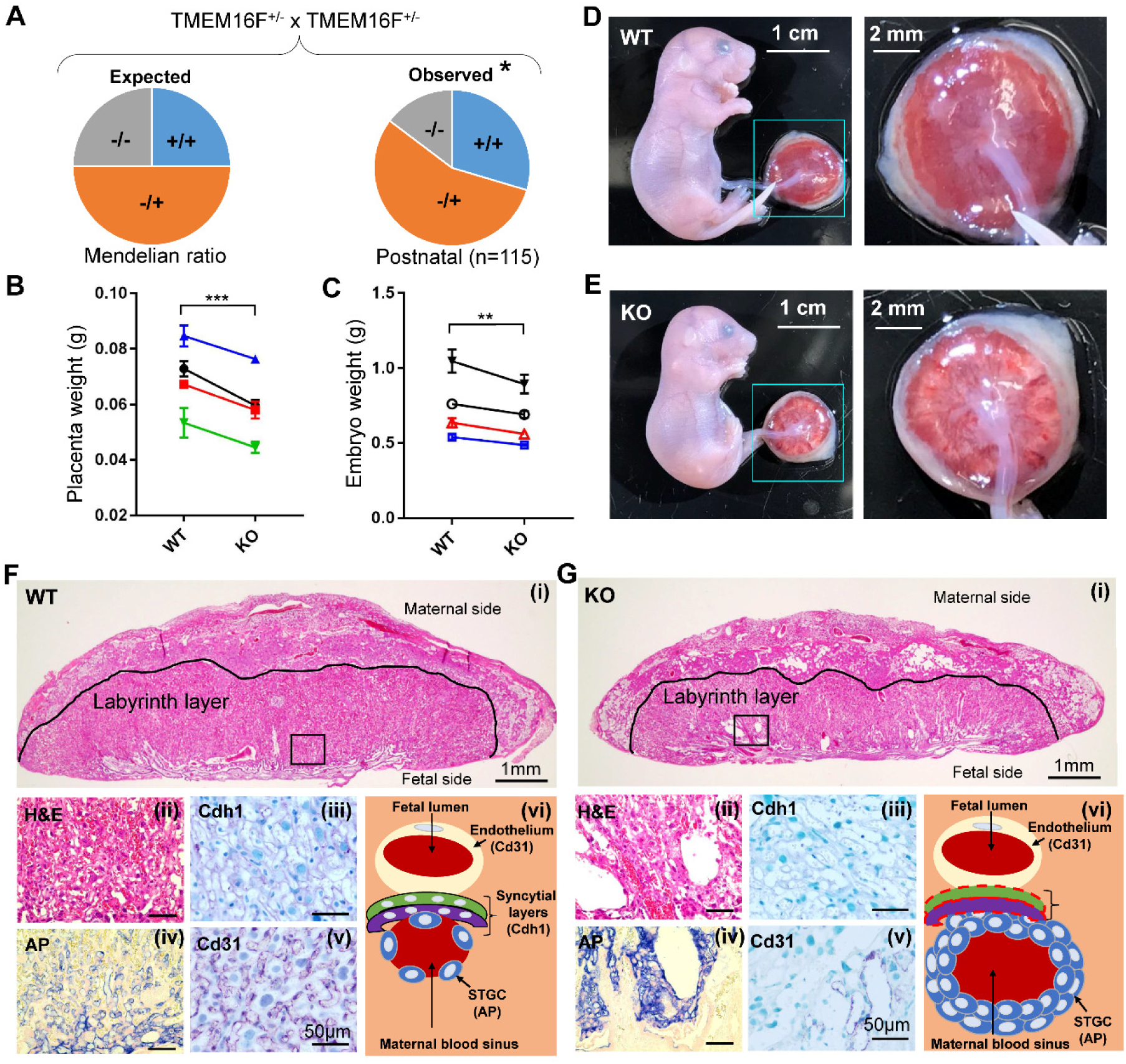
TMEM16F knockout (KO) mice exhibit perinatal lethality and defects on placenta development. (A) Significant reduction of TMEM16F^-/-^mice from the expected Mendelian inheritance. *p < 0.05 with Chi-quare test. (B, C) TMEM16F^-/-^(target deletion) mice show markedly decreased placenta weight (B) and embryo weight (C). Each data point represents the averages of the littermates with the same genotype and each line links the same litter. Two-way ANOVA. ***p < 0.001, **p < 0.01. All data represent means ± SEM. (D, E) Representative embryos and placentas from the wild-type (WT, D) and TMEM16F KO (E) mice at E18.5. The right panels show higher magnifications of the placentas with the fetal side facing up. (F, G) H&E staining (i and ii), immunostaining against E-cadherin (Cdh1, iii), alkaline phosphatase (AP, iv) and immunostaining against Cd31 (v) of the TMEM16F WT (F) and KO (G) placentas at E18.5. Cdh1, AP and Cd31 staining label syncytiotrophoblast layers, sinusoidal trophoblast giant cells (STGCs) and fetal blood capillaries, respectively. Schematics of the fetomaternal exchange anatomy from TMEM16F WT and KO mouse placentas are shown in F(vi) and G(vi), respectively. Note that Cdh1 staining of syncytiotrophoblasts in the KO labyrinth layer is greatly diminished (G(iii) and represented as red dotted lines in G(vi)). Multiple layers of sinusoidal trophoblast giant cells are densely packed together surrounding the enlarge maternal blood sinuses in the region of the KO labyrinth near the fetal interface. Cd31 staining of fetal blood vessels is markedly decreased in the KO labyrinth layer.

In addition to the enlarged maternal blood sinuses, E-cadherin (Cdh1), a marker of syncytiotrophoblasts^36^, has markedly diminished expression in the TMEM16F^-/-^ labyrinth layer (Figures 4F(iii), 4G(iii) and S4A), suggesting that the TMEM16F KO placentas have defective syncytialization. Interestingly, AP staining of the STGCs is prominently enhanced in the KO placentas compared to the WT placentas (Figure 4F(iv) and 4G(iv)). In fact, multiple layers of STGCs are observed surrounding the enlarged maternal blood sinuses whereas the WT maternal blood sinuses are usually enclosed by a single layer of STGCs. As STGCs and two layers of syncytiotrophoblasts collectively form the three-layer barriers between maternal blood and fetal vessels in the mouse labyrinth layer^37^ (Figure 4F(vi)), the observed reduction of E-cadherin staining, enhanced AP staining and increased number of STGCs surrounding the maternal blood sinuses suggest that the barrier function of the TMEM16F KO labyrinth layer is compromised likely due to deficiency in trophoblast syncytialization (Fig. 4G(vi)). TMEM16F KO STGCs likely become hyper-proliferative and form multi-layer structures to compensate the barrier defects due to deficiency in the syncytial layers and maternofetal exchange.

Consistent with the deficiency in maternofetal exchange, CD31 immunostaining of the fetal blood vessel clearly demonstrates that the fetal vessels in the KO placentas fail to develop and branch properly (Figures 4F(v), 4G(v), and S4B). Thus, the defective syncytialization and maternofetal exchange (Fig. 4G(vi)) might stress the KO placentas, which eventually causes fetal vascularization defects (Figures 4G(v) and S4B), placental insufficiency (Figure 4E and 4G) and perinatal lethality (Figure 4A and Table S1).

Similar placental defects were also observed in another TMEM16F KO mouse line generated using gene trap (Figure S5 and Table S1). The lethality of this KO line is even more prominent (Table S1) with a 4.8% postnatal survival rate in homozygotes, which is similar to the lethality observed in another TMEM16F KO mouse line (7.4% postnatal survival in homozygotes)^38^. Taking the *in vitro* and *in vivo* studies together, we therefore conclude that the TMEM16F CaPLSase plays an essential role in mediating trophoblasts PS exposure and controlling trophoblast fusion and placental development.

## Discussion

As the most studied CaPLSase, TMEM16F has been shown to play critical roles in blood coagulation^20, 21^, bone mineralization^38^, immune response^39^ and HIV infection^40^. Here we show that TMEM16F also plays an indispensable role in trophoblast fusion and placental development. When TMEM16F is genetically ablated and CaPLSase-mediated PS exposure is eliminated, BeWo trophoblasts can no longer fuse (Figure 3C) even if syncytin proteins, the key fusogens in trophoblast^12, 13^, are present. Heterologous expression of mouse TMEM16F into TMEM16F knockout BeWo cells reintroduces CaPLSase activity and cell-cell fusion (Figure 3D and 3E). Furthermore, the TMEM16F knockout mouse lines show impaired syncytiotrophoblast formation, abnormally enlarged maternal blood sinuses in the labyrinth layer and malformation of fetal blood vessels, which collectively cause perinatal lethality (Figures 4 and S4). The *in vivo* findings thus further support the essential role of TMEM16F CaPLSase in trophoblast fusion, deficiency of which will lead to defects in placental development and partial perinatal lethality to the knockout mice.

Contrasting the prevailing view that PS exposure during cell-cell fusion is a consequence of apoptosis and caspase activation^17-19^, our experiments clearly demonstrate that apoptosis is not required for CaPLSase-mediated cell-cell fusion. Application of Q-VD, a pan-caspase inhibitor, has no effects on trophoblast fusion (Figure 1D and 1E), suggesting that apoptosis and caspase-activated lipid scramblases play a dispensable role in PS exposure during trophoblast fusion. Considering the facts that 1) multinucleated syncytial cells need to stay healthy to fulfill critical physiological functions, 2) one of the advantages of cell-cell fusion is to supply the fused cells with additional genetic and metabolic materials, 3) not all fused cells have the robust phagocytic machinery as phagocytes to handle apoptotic cells, cell fusion with apoptotic cells would increase the burden of the fused cells rather than benefit their functions. With more in-depth understanding of TMEM16 CaPLSases^20^ and caspase-activated lipid scramblases^27^, we now understand that a viable cell with high CaPLSase expression can effectively externalize PS to cell surface^41^ without triggering apoptosis^30, 42, 43^. Our findings thus support that under physiological conditions, cell-cell fusion preferentially happen between healthy cells instead of involving apoptotic cells.

Cell-cell fusion is a highly coordinated, tightly regulated, multi-step process, which requires synergistic actions of multiple fusion machineries^1-3, 9^. The known mammalian fusogens show clear cell-type specific distribution^1-3^. In the placenta, syncytins are believed to interact with their receptors to promote trophoblast fusion^12, 13, 44^. Considering the broad expression of TMEM16 scramblases^45^, CaPLSase-mediated PS exposure thus may serve as a universal prerequisite for cell-cell fusion. CaPLSase-mediated PS exposure mightwork synergistically with cell-type specific fusogens to facilitate cell-cell fusion (Figure 3F). The exposed PS on cell surface may prime the cell-cell fusion sites and create an environment to enable fusogens and their partners to form stable interactions, which eventually lead to fusion pore formation. On the other hand, it is also likely that CaPLSase-mediated PS surface exposure in one fusing cell recruits some cell-surface PS receptors from another fusing cell. The interaction between PS and PS receptors, similar to the interaction between PS on the apoptotic cell surface and PS receptors in phagocytes during phagocytosis^14, 16, 46^, provides a simple and universal phospholipid-protein interaction that promotes membrane-membrane contact and fusion of various cell types. Therefore, cell-type specific fusogens and their partners might work in concert with CaPLSases to render cell-cell fusion high efficiency and high specificity. The existence of multiple, highly coordinated yet seemingly redundant cell-cell fusion mechanisms thus can provide evolutionary advantages for the reproduction and survival of an organism. All these hypotheses need to be tested in future studies.

CaPLSase-mediated cell-cell fusion mechanism will have broad implications in health and disease. In the placenta, we show that TMEM16F CaPLSase plays a critical role in placental and embryonic development (Figure 4). The TMEM16F knockout mouse placentas exhibit apparent morphological defects during development, which lead to deficient feto-maternal exchange and partial perinatal lethality. The defects in the syncytiotrophoblast layers in the labyrinth seems to be partially compensated by the hyper-proliferation of the STGCs surrounding the enlarged maternal blood sinuses (Figure 4G). The same compensatory effect was not reported in the characterizations of the Syncytin knockout mice^47, 48^. This might be the reason why the TMEM16F knockout lethal phenotype seems less severe than the Syncytin-A knockout mice. In addition, CaPLSase-mediated cell-cell fusion mechanism can also shine new lights on understanding the pathophysiology of placental disorders such as preeclampsia, pregnancy insufficiency, fetal growth retardation and preterm birth. As a general cell fusion mechanism, CaPLSase-mediated cell-cell fusion might also help understand other cell fusion events, including skeletal muscle development and repair, fertilization, osteoclast formation, cancer cell fusion and viral infection^1, 3, 14, 19, 40^. Our findings thus open new avenues of investigation to understand cell-cell fusion and membrane-membrane interactions in wide range of biological contexts.

## Acknowledgments

We are grateful to Lily Y. Jan for providing the TMEM16F deficient mice, So-Young Kim for the assistance on generating the CRISPR BeWo TMEM16F knockout cell lines, and Mathieu Flamand and Kate Meyer for helping with lentivirus generation. We appreciate Mana Parast and Jeff Everitt for their help with placenta histology, Son Le for molecular biology, and Charles Lockwood, Vann Bennet, Michael Boyce, Zhushan Zhang, Ping Dong and Son Le for their constructive comments on the manuscript. This work was supported by the National Institutes of Health (NIH-DP2GM126898 to H.Y.), the Whitehead Foundation (H.Y.), and the USF Nexus Initiative (H.P.).

## Author Contributions

H.Y. conceived and supervised the project. Y.Z. and H.Y. designed the research and performed cell fusion experiments. Y.Z. characterized the knockout mice. T.L. selected and characterized TMEM16F knockout cells and worked on all molecular biology. T.L. Y.Z. and H.Y. developed the lipid scrambling assay. Z.M. and L.P. provided human placenta tissue and performed PCR. R.G., H.P., H.K. and D.R.N. performed histology. H.Y. and Y.Z. wrote the manuscript with inputs from L.F., D.R.N, and H.P.

## Methods

### Cell lines

The BeWo cell line was a gift from Dr. Sallie Permar at Duke University. The cell line was authenticated by Duke University DNA Analysis Facility. BeWo cells were cultured in Dulbecco’s Modified Eagle Medium-Hams F12 (DMEM/F12) medium (Gibco, REF 11320-033), supplemented with 10% FBS (Sigma, cat. F2442) and 1% penicillin/streptomycin (Gibco, REF 15-140-122). All cells were cultured in a humidified atmosphere with 5% CO_2_ at 37°C.

### Human placenta tissue and primary cultured human trophoblast cells

Placenta tissues were collected under the Institutional Review Board approval (IRB# PRO00014627 and XHEC-C-2018-089) with the waiver of consent to obtain deidentified tissue that was not to be used for clinical purposes. Tissue was transported to the laboratory in Dulbecco’s Modified Eagle Medium-Hams F12 (DMEM/F12) medium (Gibco, REF 11320-033). For primary trophoblast culture, the placentas were collected from women who underwent planned cesarean delivery at term (39-40 gestational week), without labor and current or previous pregnancy complications. Placental cytotrophoblast cells were prepared using a modified method of Kliman as described previously ^49^. In brief, villous tissues were removed randomly from the maternal side of the placenta. The isolated tissues were minced after washing with normal saline and digested with 0.125% trypsin (Sigma) and 0.03% DNase I (Sigma, cat. 11284932001) in Dulbecco’s Modified Eagle Medium (DMEM) (Gibco, REF 11995-065) and incubated for 30 minutes at 37°C with agitation. The supernatant from the first digestion was discarded. Three more rounds of digestions were followed. For each digestion, the dispersed placental cells were filtered through 70 μm cell strainer (BD Falcon, cat. 352350) and centrifuged at 1000 rpm for 10 minutes. The pooled cells were then purified using a 5–50% Percoll (Sigma, cat. P1644) gradient at stepwise increments of 5%. The cytotrophoblasts between densities of 1.049 g/mL and 1.062 g/mL were collected and plated in fibronectin (10 μg/mL)-coated cover glass for culture at 37°C in 5% CO_2_-95% air in DMEM containing 10% fetal calf serum (Sigma, cat. F2442) and 1% antibiotics (Gibco, REF 15-140-122).

### Mice

The gene targeting knockout line was a kind gift from Dr. Lily Jan and has been reported previously ^21^. The heterozygous (het) mice have been backcrossed 10 generations into a C57BL6 strain. The het x het crosses generated TMEM16F^-/-^knockout mice. The gene trap TMEM16F/Ano6 knockout line was purchased from the Jackson Laboratory (C57BL/6-*Ano6*^*Gt(EUCJ0166e09)Hmgu*^, #024798). PCR genotyping was performed using tail DNA extraction. Mouse handling and usage were carried out in a strict compliance with protocols approved by the Institutional Animal Care and Use Committee at Duke University, in accordance with National Institute of Health guideline.

### TMEM16F knockout cell lines

Stable Cas9-expressing cells were generated by transducing BeWo cells with lentiCAS9-blast (Addgene, cat. 52962). To generate TMEM16F/ANO6 knockout cells, we designed sgRNAs targeting exon 2 using CHOPCHOP (http://chopchop.cbu.uib.no/index.php) and cloned these sequences into lentiguide-puro (Addgene, cat. 52962). All lentiviruses were prepared by cotransfecting HEK293T cells with the lentivector, psPAX2 and pMD2.g using TransIT-LT1 (Mirus, cat. MIR 2304). All transductions were done at a multiplicity of infection (MOI) of <1 in the presence of 4 µg/ml polybrene. Twenty-four hours post infection, cells were selected with 10 µg/ml blasticidin or 2 µg/ml puromycin for 48-72 hours and then expanded. Genomic DNA was harvested from cells one week after transduction with sgRNAs. The TMEM16F exon 2 locus was PCR amplified from genomic DNA and used in Surveyor assays (Integrated DNA Technologies, cat. 706020) to confirm the presence of insertions and deletions.

The BeWo cell populations were selected for single-cell colonies by serial dilution into 96-well plates. The cells were grown for 10-14 days, and then expanded to larger culture plates. Western blots, immunofluorescent staining and phospholipid scrambling assay were used to screen the single-cell colonies that have no TMEM16F expression and CaPLSase function. Sequences used: sgRNAs used to generate TMEM16F-KO Bewo are AATAGTACTCACAAACTCCG and TTTCCAGTGATCCCAAATCG. TMEM16F forward and reverse primers used in Surveyor assays are TTTTCAGTGGTAGACCTTGCCT and AAGTTCAGCAACCTATTCCCAA, respectively.

### Expression of murine TMEM16F in TMEM16F-KO BeWo cells

The murine TMEM16F with a C-terminal mCherry tag was transduced into TMEM16F deficient Bewo cells using lentivirus. The lentivirus was produced by co-transfecting the packaging plasmid pCMV-deltaR8.9, envelope plasmid pCMV-VSVg, and ANO6-pLVX-mCherry-C1 vector (Addgene cat. 62554) into HEK 293T cells using 1mg/mL of PEI Max 40000 (Polysciences, cat. 24765-1). After 48 hours of transfection, the lentivirus was concentrated by ultra-centrifugating the transfected HEK293T culture supernatant at 19700 rpm for 2 h at 17°C. The concentrated virus was used to infect TMEM16F-KO Bewo cells at MOI of 4 in the presence of 2 µg/mL polybrene. TMEM16F expression was detected by the mCherry signal, and the medium was replaced with the fresh medium after 48h of infection. The transduced cells were expanded 72h post transduction.

### Fluorescence imaging of Ca^2+^ and scramblase-mediated PS Exposure

To monitor Ca^2+^ dynamics, cells were incubated with 1 μM Calbryte-590 AM Ca^2+^ dye (AAT Bioquest, cat. 20701) or Calbryte-520 AM (AAT Bioquest, cat. 20650) for 50 minutes at 37 °C. After loading, the cells were washed with HBSS solution twice at room temperature to remove excessive Calbryte dyes. The Calbryte-590 loaded cells were then incubated in HBSS buffer containing 0.5 µg/mL CF 488-tagged-Annexin-V (Biotium, cat. 29011), a fluorescence PS probe. 1 μM ionomycin or 20 nM TRPV4 agonist was added to increase intracellular Ca^2+^. A Zeiss 780 inverted confocal microscope was used to simultaneously image Ca^2+^ dynamics and CaPLSase-mediated PS exposure at a 5s interval. ZEN and MATLAB were used for imaging analysis.

### Quantitative RT-PCR

Total RNA was extracted using RNeasy Mini Kit (Qiagen, cat. 74104) and quantified using NanoDrop Spectrophotometer (NanoDrop, Wilmington, DE). Total RNA (1µg) was used to generate cDNA using SuperScript III and Oligo dT (Thermo Fisher Scientific, cat. 18080051) following the manufacturer’s protocols. cDNA (25 ng) were used for qRT-PCR using 2 x IQ SYBR Green supermix cocktail (Bio-Rad, cat. 1708880). Primers used for all genes are listed in the following Table. Primers used for internal control gene glyceraldehyde-3-phosphate dehydrogenase (GAPDH) were forward (CATGAGAAGTATGACAACAGCCT) and reverse (AGTCCTTCCACGATACCAAAGT). The iCycler was programmed for an initial denaturation step of 95°C for 2 minutes, followed by a two-step amplification phase of 35 cycles of 95°C for 30 seconds, and 60°C for 1 minute while sampling for fluorescein emission. Samples were run in duplicates, and the mean cycle threshold (Ct) was normalized to the average GAPDH Ct. Fold changes were calculated using the ΔΔCt method after normalization (all gene expressions were compared with Syncytin-1).

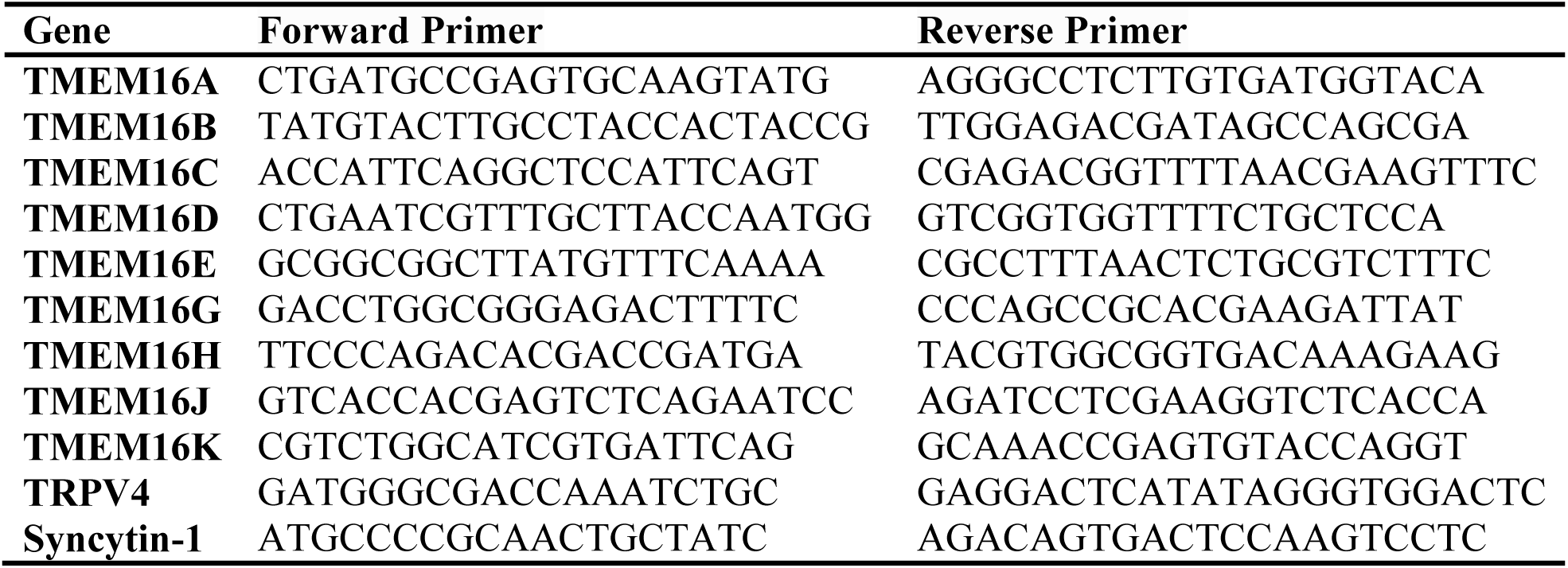

### Immunofluorescence staining

Cells grown in monolayer cultures were fixed with 4% paraformaldehyde in phosphate-buffered saline (PBS), permeabilized with 0.2% Triton X-100, and blocked with 10% goat serum prior to antibody staining. Specific primary antibodies were added at 1:500 dilution overnight. Fluorescent staining was developed using the Alexa-488 or Alexa 594 fluorescence system (Molecular Probes, cat. 35552). Cover slips were mounted onto slides with Vectashield mounting medium with DAPI (Vector Laboratories Inc, cat. H-1200). Fluorescent images were collected by using Zeiss 780 inverted confocal microscope. ZEN and ImageJ were used for imaging analysis.

### Immunoblotting

After the cells reached 80-90% confluence in 96-mm culture disk, trypsinization was used to collect the cells, which were then pelleted by centifugating at 900 rpm, 4°C. After washing twice with cold PBS, the cell pellet was lysed with RIPA lysate buffer (Thermo, cat. 89900) in the presence of 5 mM EDTA (Thermo, cat. 1861274) and 1X Protease inhibitor cocktail (Thermo, cat. 1862209). After incubating on ice for 30 mins, the lysate was centrifuged at 11000 rpm for 20-min at 4°C. The supernatant was collected and incubated with 1X Laemmli (Bio-Rad, cat. 161-0747) supplemented with 112 mM DTT (Bio-Rad, cat. 161-0611) for 30mins at room temperature, and then separated on SDS-PAGE gel. After that, Trans-Blot Turbo Transfer system (Bio-Rad) was used to transfer the proteins from the SDS-PAGE gel to PVDF membrane. The membrane was incubated in Ponceau-S stain for 15 minutes. After three-time washing with 5% acetic acids, the Ponceau-S stained membrane was imaged using ChemiDoc XRS+ System (Bio-Rad). 5% non-fat milk, 0.1% Tween-20 (Sigma, cat. P9416) was then used to block the membrane for 1-hour at room temperature. The membrane was incubated with anti-TMEM16F antibody (Millipore-Sigma, cat. HPA038958) with gentle shaking overnight at 4°C. Following three-time PBST (PBS supplemented with 0.1% Tween-20) washes, the membrane was then incubated with corresponding secondary antibody for 1-hour at room temperature. Clarity™ Western ECL Substrate kit (Bio-Rad. Cat. 170-5060) was used to activate chemiluminescent signals from protein bands. The chemiluminescent signals were detected and analyzed by ChemiDoc XRS+ System (Bio-Rad) and Bio-Rad Imaging software. The TMEM16F bands intensity from the chemiluminescence were normalized to the total protein loading detected by Ponceau-S.

### Cell-cell fusion and fusion index quantification

Syncytialization of BeWo cells was induced with forskolin and fusion index was measure using a simple live cell imaging method^24^. Briefly, BeWo cells were seeded on Poly-L-lysine (Sigma, cat. P2636) coated NO.0 cover glass for 24 hours at 5% CO_2_ at 37 °C. After 24 hours, BeWo cells were treated with 30 µM forskolin (Cell Signaling Technology, cat. 3828s) for 48 hours to induce cell fusion. Culture medium containing 30 µM forskolin was replaced every 24 h. After treatments, the cultured cells were incubated with Hoechst (Invitrogen™, cat. H3570, 1:1500) and Di-8-ANEPPS (Invitrogen™, cat. D3167, 2 µM) at 5% CO_2_ at 37 °C for 15-20 minutes, and subsequently washed in dye-free medium for twice at room temperature. Fluorescent images of six fields were randomly captured with a Zeiss 780 inverted confocal microscope using a 63x/1.4 NA Oil Plan-Apochromat DIC objective. ZEN and MATLAB were used for imaging analysis. Cells with more than 2 nuclei are defined as fused cells. To quantify cell fusion, fusion index (FI) of each sample was calculated as:

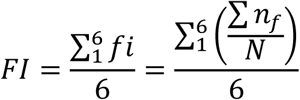

where *fi* is the fusion index of each individual random field; *n*_*f*_ is the number of nuclei from a fused cell and *N* is the total number of nuclei for each field.

To interfere with BeWo cell fusion, 0.5 μg/mL Annexin V (Biotium), 10 µM Q-VD-OPh (Sigma, cat. SML0063), or 100 nM and 500 nM GSK2193874 (Sigma, cat. SML0942) was added to culture medium 12 hour before forskolin treatment and remained throughout the experiment.

### Mouse placenta histological analysis

Mice were deeply anesthetized using isoflurane. Placentas and embryos were freshly collected and fixed in 4% paraformaldehyde (Electron Microscopy Sciences, cat. 15710) for 2 days at 4 °C and then transferred to 70% alcohol for 3-5 days before processed by using 4-hour tissue processing setting with Leica ASP6025 (Leica Biosystems). Placentas were embedded in paraffin (HistoCore Arcadia H and Arcadia C, Leica Biosystems) right after the processing and then sectioned at 5 µm by using Leica RM2255 (Leica Biosystems, Buffalo Grove, IL) for histological staining.

H&E staining was performed through deparrifinization at 60°C followed by Xylene twice (Fisher Chemical, cat. X5S-4) and a graded alcohol series 100%, 95%, 70%. Tissue was rinsed in MilliQ water for 5 minutes and then incubated in Hematoxylin 560 MX (Leica Biosystems, cat. 3801575) for 2 minutes and in running water for 1 minute. Tissue was dipped in 0.3% Ammonium Hydroxide 7 times (Fisher Chemical, cat. A669S-500) and then rinsed in running water for 1 minute. Tissue was incubated in 70% ethanol for 1 minute, followed by 10 dips in Alcoholic Eosin Y 515 (Leica Biosystems, cat. 3801615). Dehydration was performed through an alcohol gradient of 95%, 100%, and Xylene twice, 1 minute each. All slides were mounted using DPX Mounting Medium (Electron Microscopy Sciences, cat. 13512)

CD31 IHC staining was performed by using Rabbit polyclonal to CD31 (Abcam, cat. ab28364) with Avidin/Biotin Blocking Kit (Vector Laboratories, Inc., cat. SP-2001), VECTASTAIN® Elite® ABC-HRP Kit (Peroxidase, Standard) (Vector Laboratories, Inc., cat. PK-6100), and VECTOR VIP Peroxidase (HRP) Substrate Kit (Vector Laboratories, Inc., cat. SK-4600). E-Cadherin (Cdh1) IHC staining was done by using mouse monoclonal [HECD-2] to E-Cadherin (Abcam, cat. ab1416) with M.O.M. Kit (Vector Laboratories, Inc., cat. BMK2202,) incorporated with the kits used for CD31 staining. Both E-Cadherin and CD31 staining have nuclear counter staining with VECTOR Methyl Green (Vector Laboratories, Inc., cat. H-3402-500). Microscopic imaging of histology samples were taken at 20X by using an Olympus microscope (BX63l, Olympus, Center Valley, PA) with cellSens Dimension software.

Alkaline Phosphatase (AP) staining was used to assess for differentiation of sinusoidal trophoblast giant cells. Briefly, sections were de-paraffinized and rinsed in PBS, incubated in wash buffer (NTMT: 0.1 M NaCl; 0.1 M Tris, pH9.5; 0.05 M MgCl_2_; 0.1% Tween-20. Prepared fresh) and alkaline phosphatase activity was detected by using NBT/BCIP substrate to form a blue precipitate as previously described ^50^. The reaction was stopped in PBS and cells were counterstained in Nuclear Fast Red (Vector Labs, cat. H-3403) for 1 min, followed by dehydration through a graded ethanol series. Images were collected using a Leica DMR light microscope (fluorescence and bright-field) and an EVOS XL Core microscope (bright-field; Invitrogen, USA).

### Statistical analysis

The analyses of placenta and embryo weight were based on two-way ANOVA. The Chi-quare test was applied to test whether the genotyes distribution of the mice from the het x het breeding scheme obeyed the Mendelian ratios. Others statistical analyses were based on a Student’s two-tailed unpaired t-test and were performed using GraphPad Prism (GraphPad Software). Unless otherwise described, the data are representative of mean ± standard error of the mean (SEM). p values less than 0.05 were considered statistically significant.

### Data availability

All data generated or analyzed during this study are included in this published article (and its supplementary information files). Raw data and code used are available from the corresponding author upon reasonable request.

## Supplementary Figures

**Figure S1.**
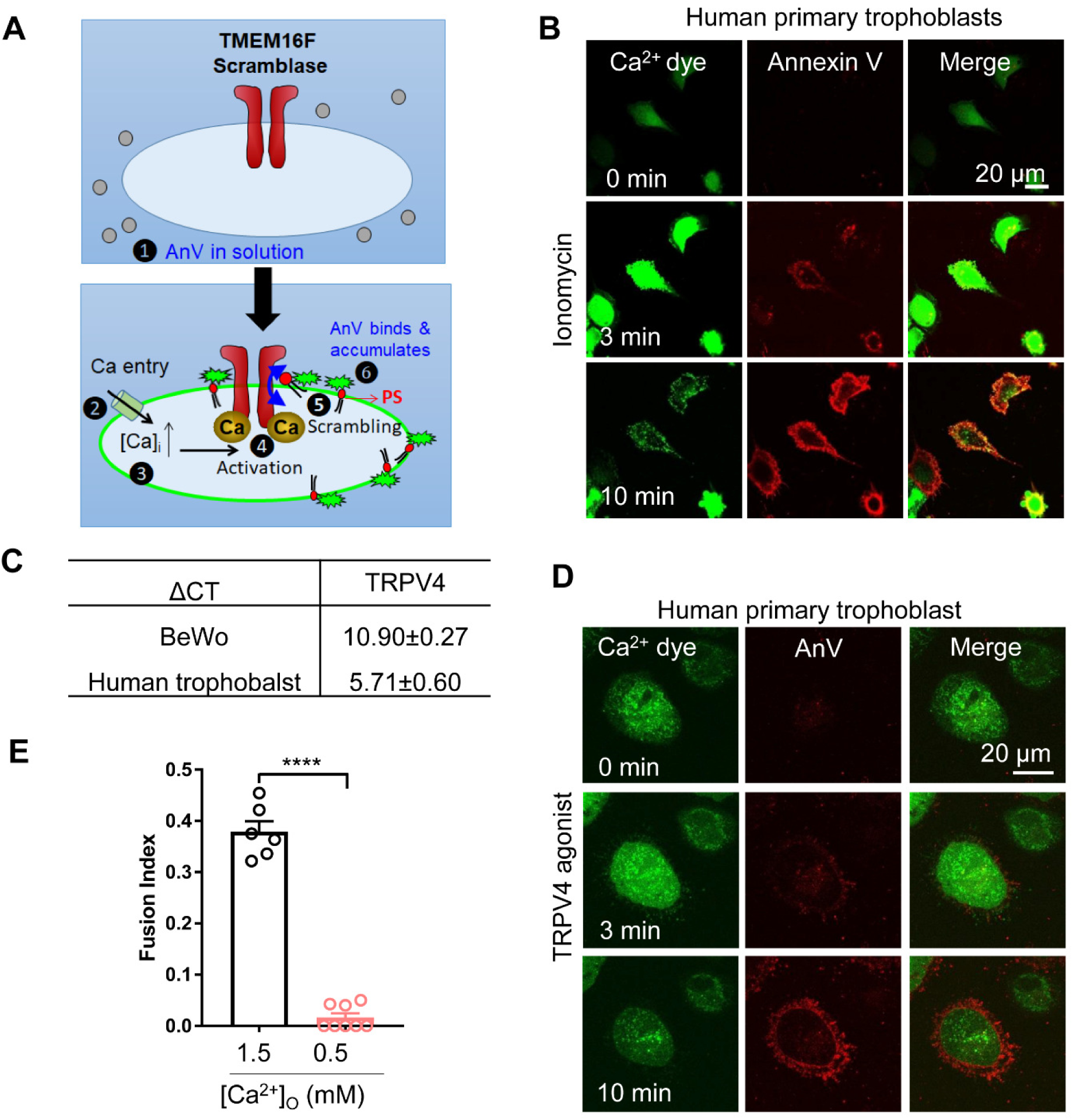
Human trophoblasts exhibit active Ca^2+^-activated phospholipid scramblase activity and Ca^2+^-dependent cell fusion. (A) An optimized microscope-based, fluorescent imaging assay to monitor Ca^2+^-activated phospholipid scramblase (CaPLSase) activities. In this assay, Ca^2+^ ionophores or agonists trigger intracellular Ca^2+^ elevation, which subsequently activates CaPLSases, resulting in PS surface exposure and cell surface accumulation of fluorescently tagged PS binding protein, Annexin V (AnV). (B) Primary human placental trophoblasts exhibit robust CaPLSase activities when triggered with 1 µM ionomycin. Ca^2+^ dye (Calbryte 520) and fluorescently tagged AnV proteins (AnV-CF594) were used to measure the dynamics of intracellular Ca^2+^ and cell surface PS, respectively. (C) qRT-PCR of TRPV4 in BeWo cells and primary human trophoblasts. ΔCT was normalized to GAPDH. (D) Primary human placental trophoblasts exhibit robust CaPLSase activities when triggered with 20 nM GSK1016790A, a potent TRPV4 channel agonist. (E) Reducing extracellular Ca^2+^ concentration ([Ca^2+^]_O_) in the cell culture medium suppressed forskolin-induced BeWo cell fusion. Each dot represents the average of fusion indexes of six random fields from one coverslip. Unpaired two-sided Student’s *t*-test. ****: p < 0.0001. Error bars indicate ±SEM. All fluorescence images are the representatives of at least three biological replicates.

**Fig. S2.**
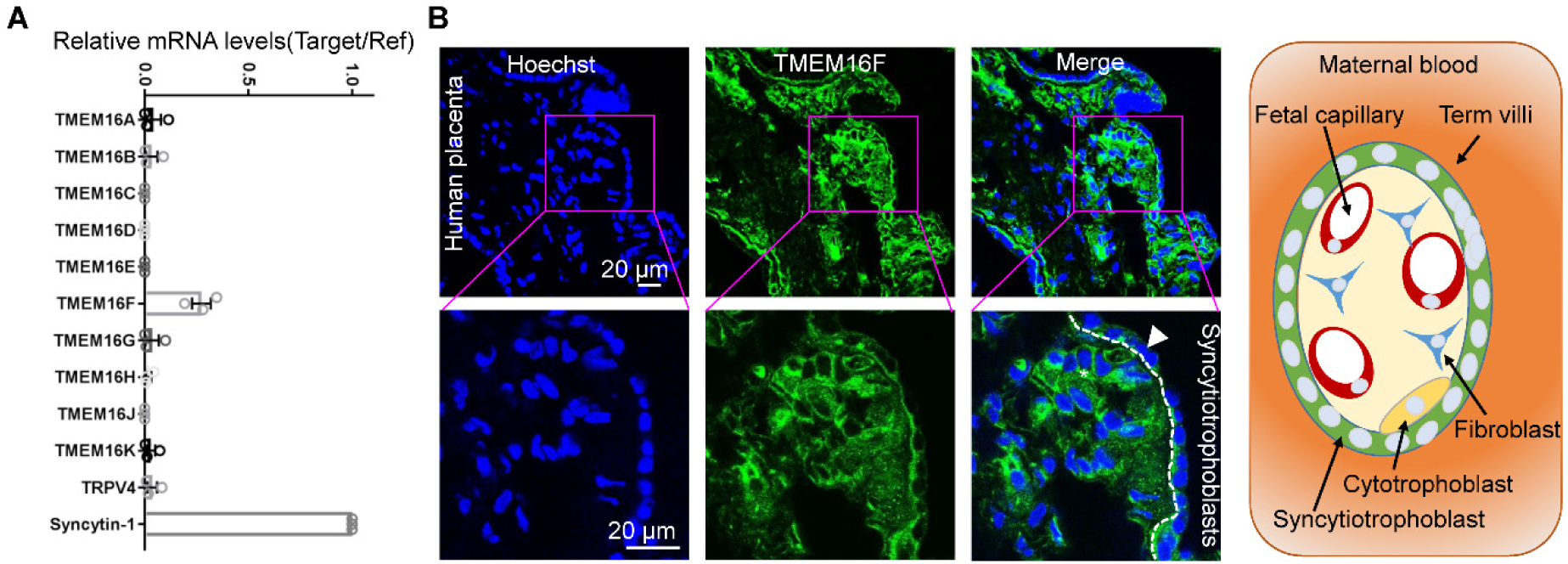
TMEM16F is highly expressed in human placental trophoblasts. (A) qRT-PCR of TMEM16 family members in primary human trophoblast shows that TMEM16F mRNA has the highest express level over other family members. All genes were normalized to GAPDH and then normalized to Syncytin-1. (B) Immunofluorescence of TMEM16F (anti-TMEM16F, green) and nuclei (Hoechst, blue) in a human term placenta sample (upper panels). Higher magnification is shown in in the lower panels. TMEM16F protein is highly expressed in the basal membrane of the syncytiotrophoblasts (arrowhead). The white dotted line demarcates the basal membrane of the syncytiotrophoblast. Schematic of the maternal-fetal interface in term placental villi is shown on the right. Images and diagram are shown in cross-sections of the villi. All fluorescence images are the representatives of at least three biological replicates.

**Fig. S3.**
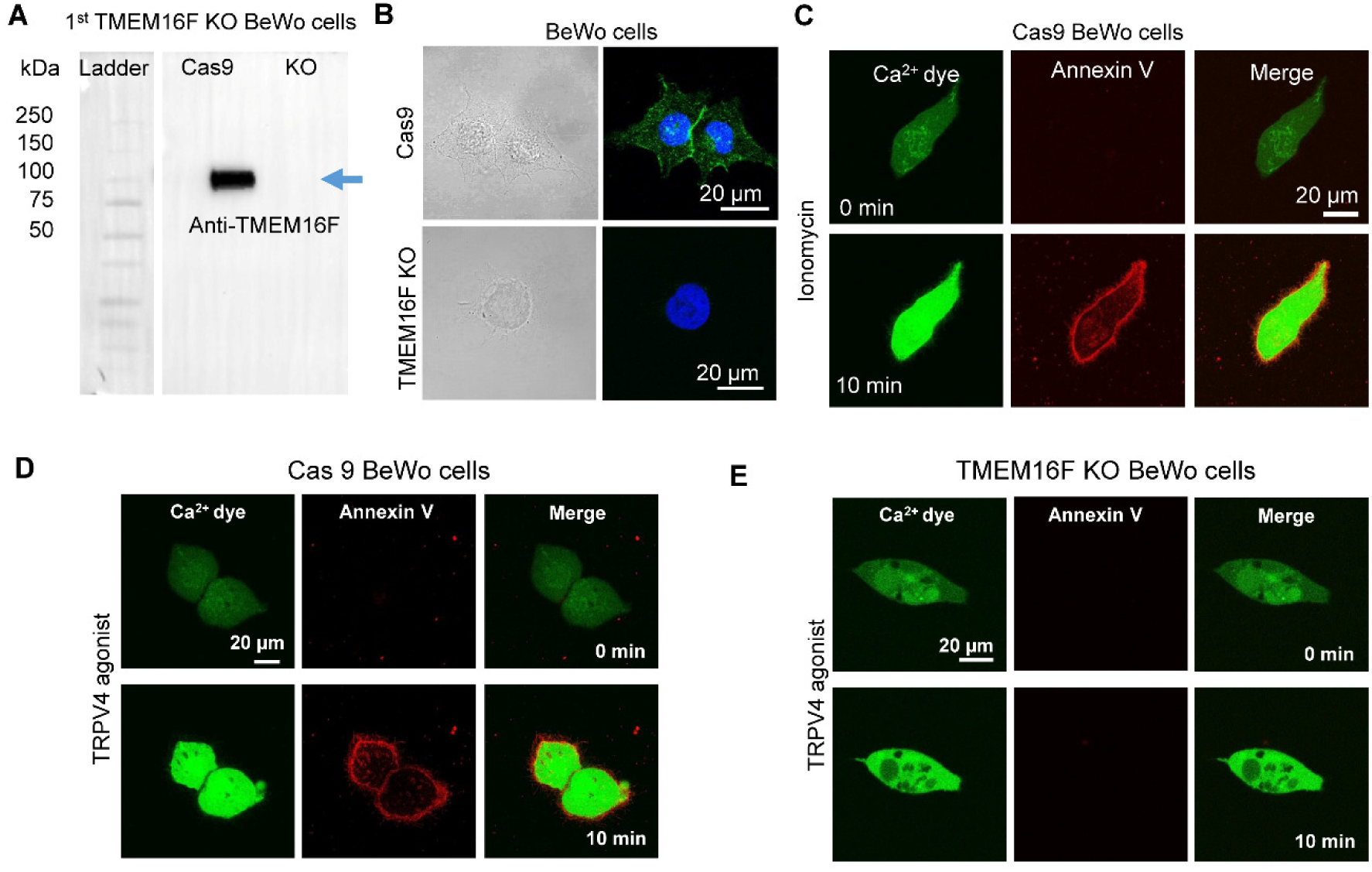
TMEM16F is responsible for the CaPLSase activity in BeWo cells. (A) Western blotting of the Cas9 control and the TMEM16F knockout (KO) BeWo cells. The arrow labels the expected TMEM16F band. (B) Immunofluorescence of TMEM16F (green) in the Cas9 control BeWo cells (upper) and the TMEM16F knockout (KO) BeWo cells generated by CRISPR-Cas9 (lower). Cell nuclei are stained with Hoechst (blue). (C) The Cas9 BeWo cells exhibit robust CaPLSase activity triggered by 1 µM ionomycin. (D) The Cas9 BeWo cells exhibit robust CaPLSase activities when triggered with 20 nM GSK1016790A, a potent TRPV4 channel agonist. (E) 20 nM GSK1016790A fails to trigger PS exposure in the TMEM16F KO BeWo cell line. All fluorescence images are the representatives of at least three biological replicates.

**Fig. S4.**
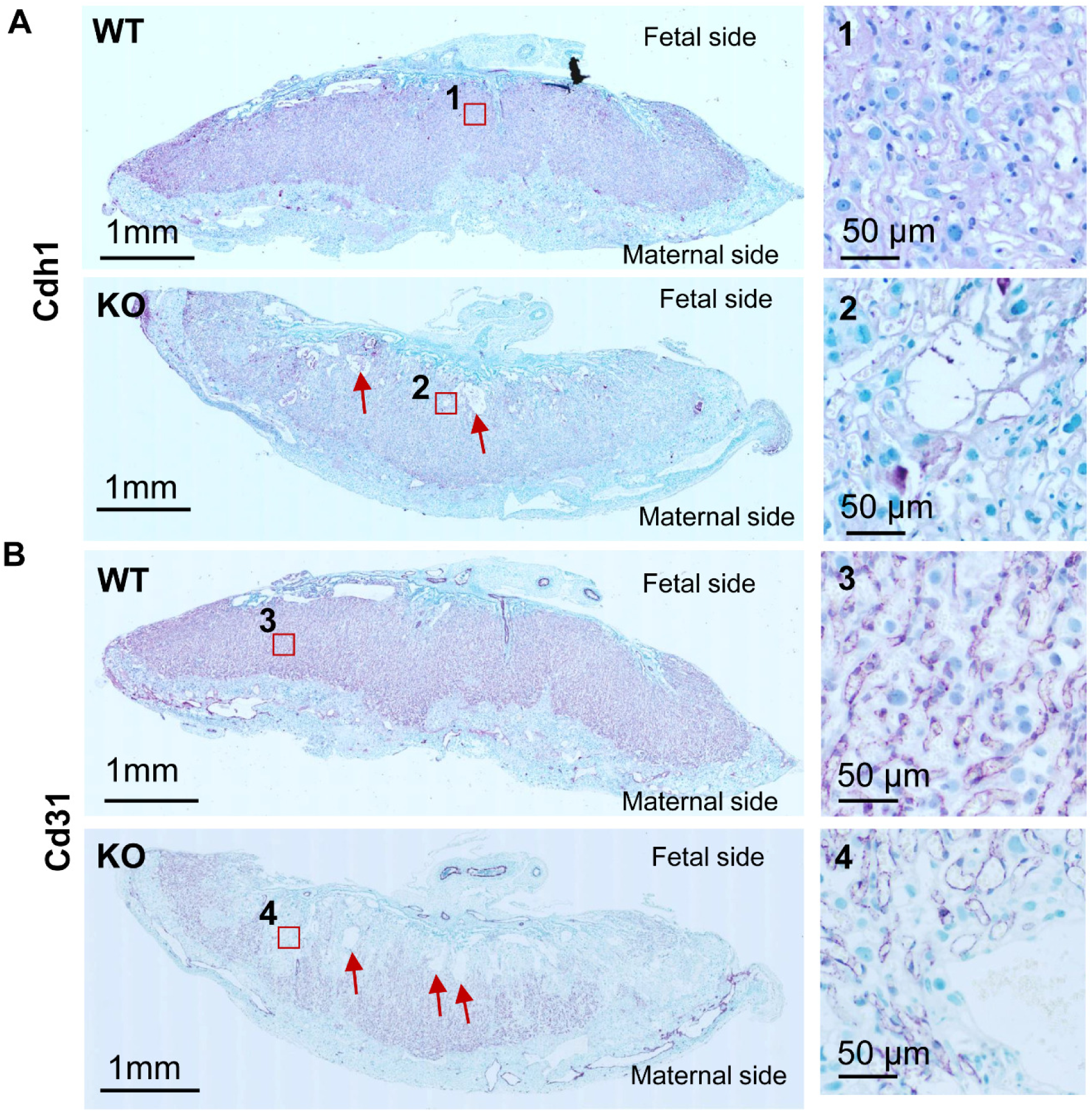
Histological changes of the placentas from the TMEM16F deficient mice. (A) Immunostaining against E-cadherin (Cdh1) on TMEM16F WT and KO placentas. The right panels show higher magnifications of the labyrinth layer of the placentas. Cdh1 staining, a syncytiotrophoblast marker, in the KO labyrinth layer is greatly diminished. Red arrows point to the enlarged maternal blood spaces of the KO placenta. (B) Immunostaining against Cd31 on TMEM16F WT and KO placentas. The right panels show higher magnifications of the labyrinth layer of the placentas. Cd31 labels fetal blood vessels and its staining is greatly diminished at the fetal interface of the KO labyrinth layer. Images are representative of at least three independent mice per genotype.

**Fig. S5.**
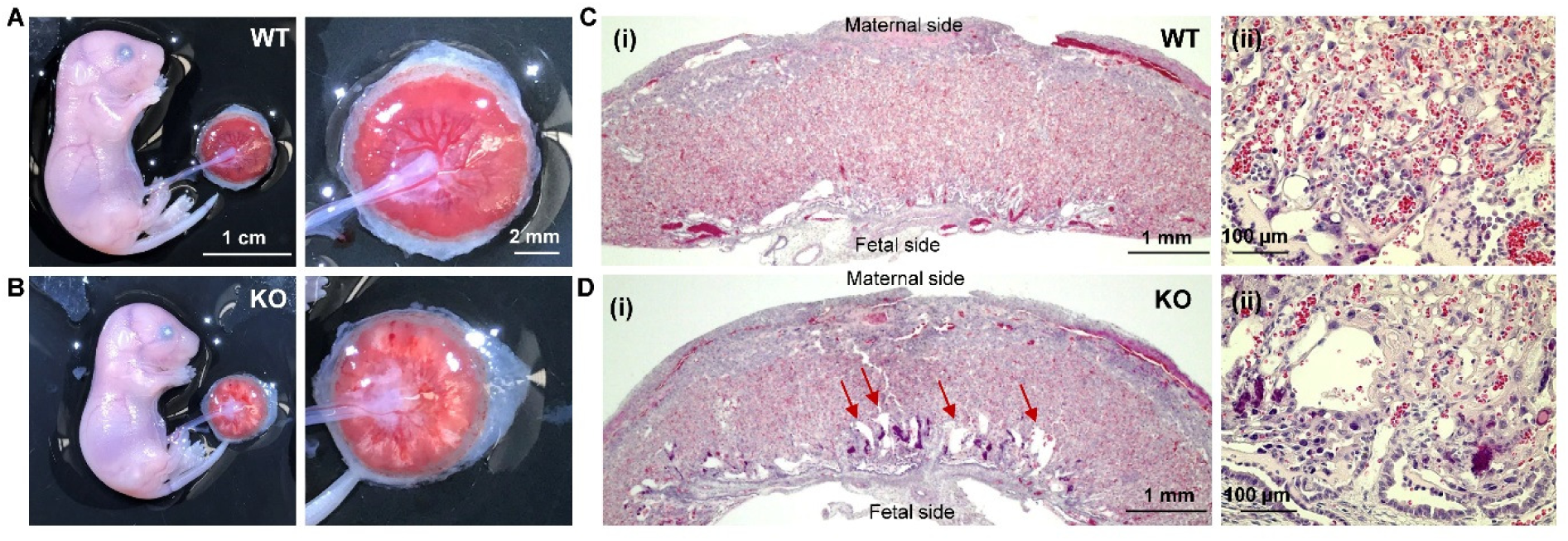
The ‘Gene Trap’ TMEM16F knockout mice exhibit similar defects on placenta morphology and development. (A-B) Representative embryos and placentas of the wild-type (WT, A) and TMEM16F deficient (KO, B) mice from the same litter at E18.5. The right panels show higher magnifications of the placentas. (C) Hematoxylin and eosin (H&E) staining of the TMEM16F WT placenta (E18.5) at low magnification (i) and the higher magnification of the labyrinth layer (ii). (D) Hematoxylin and eosin (H&E) staining of the TMEM16F KO placenta (E18.5) at low magnification (i) and the higher magnification of the labyrinth layer (ii). Red arrows point to the enlarged maternal blood spaces of the KO placenta. Images are representative of at least three independent mice per genotype.

## Supplementary Table

**Table S1.** Genotype distribution of TMEM16F knockout mice.

|  | **Total** | **WT** | **Heterozygous** | **Homozygous** |
| --- | --- | --- | --- | --- |
| Targeted Deletion* | 115 | 34 (29.6%) | 64 (55.7%) | 17 (14.7%) |
| Gene Trap**** | 104 | 31 (29.8%) | 68 (65.4%) | 5 (4.8%) |

#: het x het breeding scheme was used to generate TMEM16F knockout mice. Statistical analysis was performed using the Chi-square test. \*\*\*\*: $p < 0.0001$ . \*: $p < 0.05$ .

